# Psychological Well-being and Demographic Factors can Mediate Soundscape Pleasantness and Eventfulness: A large sample study

**DOI:** 10.1101/2020.10.16.341834

**Authors:** Mercede Erfanian, Andrew Mitchell, Francesco Aletta, Jian Kang

## Abstract

Soundscape studies aim to consider the holistic human perception of a sound environment, including both the physical phenomena and how these are mediated by internal factors: the mechanisms underpinning the interactions between these two aspects are not well understood. This study aims to assess the influence of psychological well-being and demographic factors including age, gender, occupation status, and education levels on the dimensions of the soundscape circumplex, i.e., Pleasantness and Eventfulness. Data was collected in eleven urban locations in London through a large-scale (N=1134) soundscape survey according to the ISO 12913-2 technical specifications and incorporating the WHO-5 well-being index. Linear mixed-effects modelling applying backwards-step feature selection was used to model the interactions between internal factors including psychological well-being, age, gender, occupation status, education levels and the soundscape Pleasantness and Eventfulness, while accounting for the random effects of the survey location. The findings suggest that internal factors account for approximately 1.4% of the variance for Pleasantness and 3.9% for Eventfulness, while the influence of the locations accounted for approximately 34% and 14%, respectively. Psychological well-being is positively associated with perceived Pleasantness, while there is a negative association with Eventfulness only for males. Occupation status, in particular retirement as a proxy of age and gender, was identified as a significant factor for both dimensions. These findings offer empirical grounds for developing theories of the interaction between internal factors and soundscape formation whilst highlighting the importance of the location, namely: the context.

---

Sound is a ubiquitous element in our daily lives. Despite a good deal of literature, it still strongly remains a centre of attention of many scientific communities. Looking deeper at the evolution of sound-related research in the field of engineering we see a considerable paradigm shift from noise mitigation to pleasant and restorative sound generation. This premise has been proposed with the hope to apply the existing environmental resources in order to provide a healthier and comforting acoustic environment and ultimately better quality of life (Kang, Aletta, Gjestland, Brown, Botteldooren, Schulte-Fortkamp et al., 2016; Kang, Aletta, Oberman, Erfanian, Kachlicka, Lionello et al., 2019). Hence, the soundscape concept, which places the emphasis on the human perception of the acoustic environment in context has emerged to support this premise.

Despite the strong evidence that research has brought for the soundscape, our understanding of the action of the Peripheral and Central Nervous System (PNS and CNS) associated with environmental sound interpretation and the factors influencing the perception of sound is still evolving and a matter of dispute among scientific communities. Understanding of the soundscape is intimately tied to certain key factors known as primary factors of the soundscape comprising acoustic properties (physical features) of the sound such as frequency/ pitch (Kumar, Forster, Bailey, Griffiths, 2008; Patchett, 1979) and intensity/loudness (Kaya, Huang, Elhilali, 2020) and secondary influences like emotions and personality traits (McDermott, 2012).

## Pleasantness and eventfulness as key components of soundscape

Understanding the soundscape concept and its components largely depends on understanding the circumplex model of affect, proposed by James Russell (Russell, 1980). The circumplex model delineates the entanglement of the emotions and their neural substrates, opposing the classic model of discrete basic emotions (Panksepp, 1998; Tomkins, 1962).

This model suggests that all affective states, described with descriptors such as alert, tense or serene, arise from cognitive interpretations of core physiological and neural sensations. These affective states are produced by two fundamental neurophysiological systems, including two orthogonal continuums: valence and arousal, which can be discerned as a linear combination or as fluctuating degrees of activation (Posner, Russell, Peterson, 2005).

Valence refers to whether an emotion is experienced as pleasant/positive or unpleasant/negative and is distributed horizontally on the circumplex space (on the X-axis). Arousal refers to whether an emotion is physiologically activating (high arousal; e.g., excited) or deactivating (low arousal; e.g., calm) (on the Y-axis) (Russell, 1980). High arousal is associated with activation of the sympathetic components of the Autonomic Nervous System (ANS) (e.g., increased heart rate) whereas low arousal is attributed to parasympathetic activation (e.g., slower heart rate).

Similarly, the soundscape entails two main perceptual attributes: pleasantness and eventfulness that are different from the physical properties of the acoustic environment and by which the listeners appraise the quality of sounds (International Organization of Standardization Technical Specification, 12913-3:2019) ^1^. Soundscape pleasantness refers to the emotional magnitude of the sound perception, while soundscape eventfulness is attributed to the intensity of the sound perception (Erfanian, Mitchell, Kang, Aletta, 2019). Like the Russell’s model structure, the common model of representing soundscape is a bi-dimensional circumplex model with pleasantness on the X-axis and eventfulness on the Y-axis, proposed by Axelsson, Nilsson, Berglund (2010).

In their study, three primary dimensions of soundscape perception were extracted from participants’ responses to complex sound samples measured on 116 attributes, using Principal Components Analysis. The first component was found to represent pleasantness (aligning with attributes such as comfortable, appealing, uncomfortable, disagreeable, and inviting) and explained 50% of the variance in the dataset. The second component was found to represent eventfulness (eventful, lively, uneventful, full of life, and mobile) and explained 18% of the variance. The third component was found to represent familiarity (commonplace, common, and familiar) and explained 6% of the variance. In their final model, these attributes reduced to eight primary unidimensional scales of pleasant, vibrant, eventful, chaotic, annoying, monotonous, eventful and calm and the reduced attributes collapsed into two orthogonally positioned components of pleasantness and eventfulness (See ‘Outcome variables’).

Likewise in the current study, we employed the circumplex model reported in a two-dimensional scatter plot with coordinates for the two dimensions ‘Pleasantness’ plotted on X-axis and ‘Eventfulness’ plotted on Y-axis, taking into account the features of the locations. To differentiate these complex components from classic pleasantness and eventfulness in Axelsson’s model, they will appear with the first letter capitalized throughout the text.

## Psychological well-being and soundscape

There are understudied secondary factors that may be linked to the perception of the acoustic environment, such as psychological well-being (Aletta, Oberman, Mitchell, Erfanian, Lionello, Kachlicka et al., 2019).

Individuals with an aberrant psychological state and poor mental health may experience environmental inputs differently to those people who do not experience such issues given that emotions, as one of the core components of psychological well-being, and sensory perceptions are closely intertwined (Kelley & Schmeichel, 2014). As reported in the relevant literature, the impact of psychological well-being is consistent among all perceptual modalities such as vision (Zadra & Clore, 2011), tactile (Kelley & Schmeichel, 2014), olfactory (Krusemark, Novak, Gitelman, 2013), and auditory (Riskind, Kleiman, Seifritz, Neuhoff, 2014). In parallel, studies in the field of psychopathology elucidated that individuals with poor psychological well-being, such as the clinically depressed, maintain bias and anomalous cognition, leading to inaccurate and distorted perception (Beck’s cognitive theory) (Clark & Beck, 2010).

## Demographic factors and soundscape

The perception of the acoustic environment or soundscape involves the sensation, identification, organization, and interpretation of ongoing omnipresent auditory information (Goldstein, Brockmole, 2016).

Soundscape does not always maintain consistency and shows a huge variation across individuals and on a general scale, among populations (Schneider & Wengenroth, 2009; Weinstein,1978). There is evidence to suggest that the differences in the demographic characteristics like gender (Xiao & Hilton, 2019; Gulian & Thomas 1986), age (Zhang & Kang, 2007), and educational background (Zhang & Kang, 2007) may determine the way we perceive environmental sounds. Additionally, these individual differences can potentially reflect in various perceptual properties, implying the difference between the encoding of certain auditory information between individuals such as pitch (Coffey, Colagrosso, Lehmann, Schönwiesner, Zatorre, 2016) or loudness (Berthomieu, Koehl, Paquier, 2021).

However, the results from past studies have, for a good part, remained inconclusive or inconsistent.

## The current study

Whilst previous research has substantially advanced our knowledge of the soundscape determinants, past studies results are predominantly limited, often focussing on controlled laboratory-based experiments, individuals with psychopathology (i.e., depression) and investigating simple tones rather than complex sounds (Riskind et al., 2014; Laufer, Israeli, Paz, 2016). In addition, the impact of psychological well-being in the context of the soundscape, by its current definition, has still largely been unexplored. So, our first aim is to understand if high levels of psychological well-being are associated with increased soundscape pleasantness and eventfulness.

The second aim of the study is to determine the associations between the soundscape and demographic factors, given there is insufficient consensus in the literature, studies are restricted to limited case studies (i.e., Peace Gardens in Sheffield – the UK) or a single ethnicity (i.e., Chinese) (Fang, Gao, Hedblom, Xu, Xiang, Hu, 2021; Ismail, 2014; Yang & Kang, 2005). We asked if age, gender, ethnicity, education level, and occupation are status associated with the soundscape Pleasantness and Eventfulness.

In this large-scale study, we explore the association of psychological well-being, demographic factors with soundscape among the members of the public with presumably no apparent psychopathology in an immersive environment with diverse demographic characteristics such as ethnicity (i.e., American, Italian) and occupation status (i.e., student, retired).

## Methods

The study was approved by the local ethics committee of University College London (UCL), BSEER, Institute for Environmental Design and Engineering (IEDE) (Dated 11-10- 2019).

### Participants

The present work is a large-scale study with data collected from the general members of the public in several locations in London with varying acoustic features. All passers-by at the data collection locations were approached in 11 locations/sites in London by the researchers and were asked if they were willing to participate in the study. Locations were selected which represented a variety of usage types, visual character, and acoustic characteristics. The minimum and maximum value of several acoustic metrics recorded at each location during the survey sessions are presented in Table B.1 in Appendix B. Only individuals on the phone, with headphones on due to attention distraction, or individuals that were deemed to be younger than 18 years old (proxy consent required) were excluded from the data collection. The total number of surveys that were originally collected from the sites was 1467.

### Measures and independent variables

The questionnaire, presented in full in Appendix A, comprising 38 items, is an adapted version of ISO/TS 12913-2:2018 ^2^ Method ‘A’ (urban soundwalk method) (Axelsson, 2012; ISO, 2018) and WHO-5 well-being index (World Health Organization, 1998), as well as demographic information. In order to answer the questions raised in this study the authors only report some sections of the questionnaire which then undergo the statistical analyses.

### Perceived affective quality/Perceptual attributes

The perceived affective quality (PAQ) of the sound environment as adopted in the method ‘A’, described in the ISO/TS 12913-2:2018, consists of category scales containing five response categories, based on the Swedish Soundscape Quality Protocol (SSQP; 41) (ISO, 2018). It includes a question ‘to what extent they agree/disagree that the present surrounding sound environment is …’. The participants judged the quality of the acoustic environment by 8 adjectives: pleasant, chaotic, vibrant, uneventful, calm, annoying, eventful, or monotonous. The answers were presented in a 5-point Likert scale ranging from ‘strongly disagree = 1’ to ‘strongly agree = 5’. The perceptual attributes measure as a unidimensional measuring tool for the perception of the acoustic environment has not been validated to this date. The PAQs were utilized as aggregated values to construct the principal components of the soundscape (Pleasantness and Eventfulness) (See ‘Outcome variables’).

In order to maintain data quality and exclude cases where respondents either clearly did not understand the PAQ adjectives or intentionally misrepresented their answers, surveys for which the same response was given for every PAQ (e.g., ‘Strongly agree’ to all 8 attributes) were excluded. This is justified as no reasonable respondent who understood the questions would answer that they ‘strongly agree’ that a soundscape is pleasant and annoying, calm and chaotic, etc. Cases where respondents answered ‘Neutral’ to all PAQs are not excluded in this way, as a neutral response to all attributes is not necessarily contradictory. In addition, surveys were discarded as incomplete if more than 50% of the PAQ and sound source questions were not completed.

### Psychological well-being/WHO-5 well-being index

WHO-5 well-being index asks how individuals have been feeling over the last two weeks such as ‘I have felt cheerful and in good spirits’. WHO-5 has been designed for multiple research and clinical purposes, covering a wide range of mental health domains namely perinatal mental health, the geriatrics mental health, endocrinology, clinical psychometrics, neurology, and psychiatric disorders screening.

The WHO-5 well-being index is known to be one of the most valid generic scales for quantification of general well-being. In terms of the construct validity of the scale, WHO-5 showed to have properties that are a coherent measure of well-being (Topp, Østergaard, Søndergaard, Bech, 2015). With regards to relevant literature, WHO-5 confirmed that all items constitute an integrated scale in which items add up related information about the level of general psychological well-being among both youngsters and elderlies (Blom, Bech, Hogberg, Larsson, Serlachius, 2012; Lucas-Carrasco, Allerup, Bech, 2012). For the purpose of analysis, a composite WHO-5 score is calculated by summing the responses to each of the 5 questions (coded from 0 for at no time to 5 for all of the time), then multiplying by 4 to get a single score which 0 (the lowest level of well-being) to 100 (the highest level of well-being) (Topp et al., 2015).

### Demographic characteristics

Demographic characteristics were presented such as age, gender (male, female), education level (some high school, high school, trade/technical/vocational training, university, and postgraduate), occupational status (employed, unemployed, retired, student, employed- student, other and rather not say), and ethnicity (Asian, black/Caribbean, middle eastern, white, and mixed). Some blank spaces were provided if they wanted to add further information. At the end of the survey, participants had the opportunity to write down any additional questions or remarks and were thanked for their participation.

### Outcome variables (the soundscape Pleasantness and Eventfulness)

The soundscape data were analysed according to the procedure laid out in Part 3 of the ISO 12913 ^3^ standard series. In order to ease data analysis and modelling the standard suggests a method to collapse the perceived affective quality responses for each of the 8 down to a 2- dimensional coordinate scatter plot with continuous values for ‘Pleasantness’ on the X-axis and ‘Eventfulness’ on the Y-axis. These coordinates are then normalized to between -1 and 1 (per the recommendation of ISO/TS 12913-3:2019). These dimensions were calculated as shown in Formulas (1 & 2):

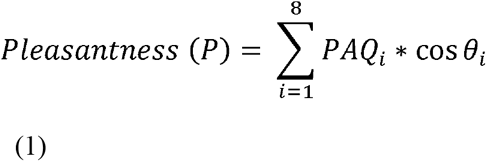

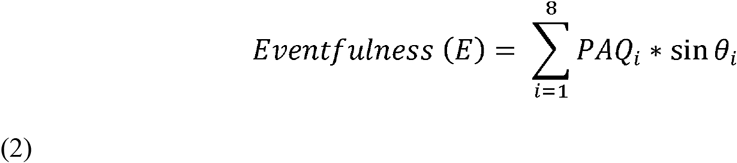

where, PAQ1 = pleasant, θ_1_ = 0°; PAQ2 = vibrant, θ_2_ = 45°; PAQ3 = eventful, θ_3_ = 90°; PAQ4 = chaotic, θ_4_ = 135°; PAQ5 = annoying, θ_5_ = 180°; PAQ6 = monotonous, θ_6_ = 225°; PAQ7 = uneventful, θ_7_ = 270°; PAQ8 = calm, θ_8_ = 315°.

### Survey procedure

The participants were approached and asked if they were interested to participate in the study. All participants received information about the aim of the study, its procedures, confidentiality of research data, and how to contact the investigators, the supervisor of the project, or a member of the ethical committee. An informed consent document was given to participants, who declared to have read and understood the general information, take part voluntarily, and have understood the fact that they can stop their participation and withdraw their consent, anytime, and without any consequences. They could start filling in the questionnaire if the participant gave his/her consent. If they had no questions, they received either a paper version or an e-version of a questionnaire via a 10-inch tablet. The online questionnaires were collected and managed using REDCap electronic data capture tools hosted at UCL (Harris, Taylor, Minor, Elliott, Fernandez, O’Neal et al., 2019) and typically took between 5 and 10 minutes to complete. The goal of the researchers on-site was to collect a minimum of one-hundred questionnaires from each selected site/location, which was typically achieved over a period of 2-3 days each consisting of approximately a 4-hour session. In some cases, either due to extenuating circumstances, time constraints, or excluded surveys, the full one hundred surveys were not achieved. The data was collected from 28^th^ February 2019 to 18^th^ October 2019 between 11 am to 3 pm.

During the survey period, acoustic and environmental metrics were simultaneously collected through binaural recordings, a calibrated sound level meter (SLM), and an environmental meter collected temperature, lighting level, and humidity data. The SLM was set up in the space in which the questionnaires were conducted and left running for the full duration of the survey in order to characterize the acoustic environment. The environmental metrics were not reported in this study since they were not in the scope of this paper but are included in the Appendices in order to provide context for the interested readers. The full protocol and data treatment as part of the SSID Database creation are described in detail by Mitchell and colleagues (Mitchell, Oberman, Aletta, Erfanian, Kachlicka, Lionello et al., 2020).

### Data analytic analysis strategy

#### Missing data, checking for outliers and data scaling

Prior to the data analysis, we imputed missing data and the imputed data was used across all analyses. Missing education values were imputed with the mode value (university). Missing values for age were imputed with the median age value (29). WHO-5 (psychological well-being) missing values were imputed with the median value (64). We excluded those who responded non-conforming (N=4) or decline (N=21) (with no response) for gender, due to the very small sample size and to simplify the effects of gender (initial number of collected data = 1467, data included in the analysis = 1134).

We took a lenient approach to outliers. Due to the nature of survey data, it was typically inappropriate to remove data solely because it represented a deviation from the typical response. However, we wanted to catch data which was incorrect, intentionally wrong, or a typo and then removed them. For the most part, this was handled with our data quality method implemented in REDCap, to ensure the SSQP/perceptual attributed values (N = 8) were filled-in such that they complied with the circumplex theory to a minimum degree. We were, therefore, only looking for values which were extreme outliers or impossible.

#### Correlation between predictors and output variables

To establish the linearity between all pairs of variables including the predictors and outcome variables, Pearson correlation coefficient, Analysis of Variance (ANOVA) and Chi-square were performed between psychological well-being, age, gender, ethnicity, education level, occupation status and the soundscape Pleasantness and Eventfulness (Table 2).

#### Model specification (linear mixed-effects modelling)

Linear mixed-effects regression (LMER) with random intercept and fixed slope, using backward stepwise feature selection was utilized to a) identify the association of our features of interest (FOIs) including psychological well-being, age, gender, education levels, ethnicity, occupation status, and their interaction terms with the soundscape Pleasantness and Eventfulness and, b) accommodate associations within participants among locations. In order to account for latent differences in the pleasantness and eventfulness ratings of various locations, the intercepts of each model are allowed to vary as a function of the location. Therefore, the model is constructed with two levels – the individual level (the random effects) and the location level (the fixed effects). Separate models were constructed for each Pleasantness and Eventfulness, and take the form (Formula 3 and 4):

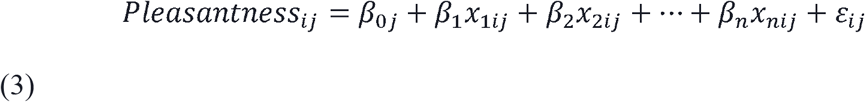

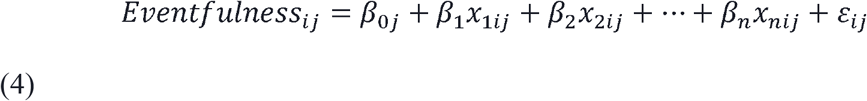

Where *Pleasantness_ij_* or *Eventfulness_ij_* are the dependent variable value for individual *i* in Location *j*; β*_0j_* is the intercept for Location *j*; β*_1_* through β*_n_* are the slopes relating the independent variables *x_1_* through *x_n_* to the dependent variable; x*_1ij_* through *x_nij_* are the dependent variables for individual *i* in Location *j*; ε*_ij_* is the random error for individual *i* in Location *j*. In turn, β*_0j_* can be expressed as:

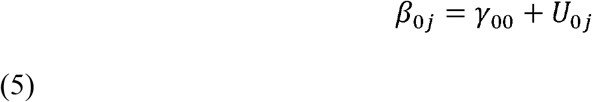

where γ*_00_* is the mean intercept across Locations; and *U_0j_* is the unique effect of Location *j* on the intercept. In a random intercept model, the slope coefficients (*β_n_*) are considered fixed across the locations (hence, labelled as the fixed effects) indicating that the relationship between the dependent variable (e.g., age, gender, etc.) and the independent variable (Pleasantness or Eventfulness) is the same for all locations, while the general Pleasantness of the location is accounted for by the varying intercept.

In order to identify the significant FOIs within the multi-level structure, we employed a stepwise feature selection on the fixed effects portion of the mixed-effects model, with an inclusion threshold of *p <* 0.05. Since this model includes only the LocationID at the random effects level, only the fixed effects are reduced in the feature selection process. To check for multicollinearity among the selected features, the variance inflation factor (VIF) was calculated and a threshold of VIF < 5 was set. Any features which remained after the backwards stepwise selection which exceeded this threshold were investigated and removed if they were highly collinear with the other features. Once the feature selection process is completed, the final model with only significant FOIs included is fit and the table of the model coefficients is printed along with plots of the random effects and z-scaled and non-standardized estimates terms.

The model fitting and feature selection was performed using ‘lme4‘ (version 1.1) and the ‘step‘ function from ‘lmerTest‘ (version 3.1.3) (Kuznetsova, Brokhoff, & Christensen, 2017) in R statistical software (version 4.0.3) (R Core Team, 2020). The summaries and plots were created using the ‘sjPlot‘ package (version 2.8.6) (Lüdecke, 2018).

## Results

The setup and procedures of this study allowed us to test a large group of participants with high diversity with rather various demographics including gender, age, education level, occupation status, and ethnicity (N= 1134) (Table 1).

**Table 1.** The sample demographic characteristics.

| Demographic characteristics | N (%) |
| --- | --- |
| N = 1134 | Age mean = 34.67 years ± 15.11 |
| Gender |  |
| Female | 610 (53.79) |
| Male | 524 (46.2) |
| Age |  |
| 18-30 | 627 (55.29) |
| 31-40 | 195 (17.19) |
| 41-50 | 112 (9.87) |
| 51-60 | 97 (8.55) |
| 61-70 | 72 (6.34) |
| 71+ | 31 (2.73) |
| Education Level |  |
| Some high school | 22 (1.2) |
| High school graduate | 315 (17.3) |
| Trade/ technical/ vocational training | 51 (2.8) |
| University (undergraduate/bachelor) | 422 (32.1) |
| Postgraduate degree (master) | 324 (17.8) |
| Occupation Status |  |
| Employed | 613 (54.05) |
| Unemployed | 25 (2.2) |
| Retired | 84 (7.4) |
| Student | 348 (30.6) |
| Employed-Student | 5 (0.4) |
| Other | 44 (3.8) |
| Rather not say | 15 (1.3) |
| Ethnicity |  |
| White | 806 (44.2) |
| Mixed/Multiple ethnic groups | 63 (3.5) |
| Asian/Asian British | 156 (8.6) |
| Black/African/Caribbean/Black British | 31 (1.7) |
| Middle Eastern | 23 (1.3) |
| Rather not say | 55 (3) |

### Correlations

The correlation matrix for all study measures is demonstrated in Table 2. Age was negatively correlated with Eventfulness, whereas it was positively correlated with Pleasantness. Gender appeared to be independent of Eventfulness but positively correlated with Pleasantness. Education was positively correlated with both Pleasantness and Eventfulness. Whilst psychological well-being exhibited positive and statistically significant correlations with Pleasantness, it was negatively correlated with Eventfulness. It is worth noting that occupation is significantly correlated with all other independent variables considered in the study and highly correlated with age, although it is not significantly correlated with either of dependant variables.

**Table 2.** Correlation coefficients for study variables.

| Factors | Age | Education | Ethnicity | Eventful | Gender | Occupation | Pleasant |
| --- | --- | --- | --- | --- | --- | --- | --- |
| Age |  |  |  |  |  |  |  |
| Education | 0.32 |  |  |  |  |  |  |
| Ethnicity | 0.23 | 0.04 |  |  |  |  |  |
| Eventful | **-0.11\*\*\*** | **0.1\*\*** | 0.08 |  |  |  |  |
| Gender | **0.1\*\*\*** | 0.05 | **0.08\*** | 0.05 |  |  |  |
| Occupation | **0.71\*\*\*** | **0.19\*\*\*** | **0.13\*\*\*** | 0.15 | **0.1\*\*** |  |  |
| Pleasant | **0.12\*\*\*** | **0.11\*\*** | 0.09 | **-0.91\*\*\*** | **0.06\*** | 0.16 |  |
| Psychological Well-being | **0.12\*\*\*** | 0.1 | **0.1\*** | **-0.12\*\*\*** | 0.02 | 0.16 | **0.14\*\*\*** |

\*\*\* $p < 0.0005$ , \*\* $p < 0.005$ , \* $p < 0.05$

### Linear mixed-effects modelling

The linear mixed-effects regression derived regularized models of the soundscape Pleasantness and Eventfulness. This model was then reduced via backward stepwise feature selection. Table 3 presents the soundscape Pleasantness and Eventfulness models, including non-standardized and standardized estimate values and CIs for the selected features that survived from the initial model. After the feature selection, age, education, and ethnicity were not found to be significant features in either the Pleasantness or Eventfulness models. It should be noted, however, that the presence of one feature (e.g., occupation) which is highly correlated with another (e.g., age and gender) may cause one of the features to not meet the threshold of significance when both are included, causing it to be removed during the stepwise feature selection. Nonetheless, it may be that, in a final model which included either of these features (but not both), they would each be considered significant. In this way, even though occupation was selected during this process, age may also have been considered significant, when not considering occupation (See Appendix C).

**Table 3.** Fixed and random effects in a linear mixed model explaining variations in the soundscape Pleasantness and Eventfulness while controlling for psychological well-being and demographic factors. The standardized estimates are calculated by refitting the model on standardized data scaled by subtracting the mean and dividing by 1 SD, allowing a comparison of all features.

| Predictor | Pleasantness |  |  | Eventfulness |  |  |
| --- | --- | --- | --- | --- | --- | --- |
|  | Estimates | Std. Est | 95% CI | Estimates | Std. Est | 95% CI |
| Psychological Well-being | **0.001\*\*** | **0.03** | 0.01, 0.05 | 0.001 | 0.01 | -0.02, 0.04 |
| Gender (male) | - | - | - | **-0.08\*** | **-0.04** | -0.07, -0.00 |
| Occupation (Rather not say) | **-0.19\*** | **-0.19** | -0.36, -0.02 | **0.7\*\*\*** | **0.02** | -0.13, 0.17 |
| Occupation (Retired) | **0.1\*\*** | **0.10** | 0.03, 0.18 | **-0.18\*\*** | **-0.11** | -0.18, -0.04 |
| Occupation (Unemployed) | 0.01 | 0.01 | -0.13, 0.14 | **0.01\*\*** | **0.18** | 0.06, 0.3 |
| Psychological Well-being x Gender (male) | - | - | - | **-0.001\*** | **-0.04** | -0.07, -0.00 |
| Psychological Well-being x Occupation (Rather not say) | - | - | - | **-0.01\*\*\*** | **-0.21** | -0.33, -0.09 |
| **Random Effects** |  |  |  |  |  |  |
| $\sigma^2$ | 0.11 | | | 0.08 | | |
| $\tau_{00}$ | 0.06<sub>Location</sub> | | | 0.01<sub>Location</sub> | | |
| ICC | 0.35 |  |  | 0.15 |  |  |
| N | 11 |  |  | 11 |  |  |
| Observations | 1134 |  |  | 1134 |  |  |
| Marginal $R^2$/Conditional $R^2$ | 0.014/0.354 | | | 0.039/0.181 | | |
| AIC | 779.125 |  |  | 451.351 |  |  |

$p < 0.05$ , $**p < 0.01$ , $***p < 0.001$

The final models found that a higher level of psychological well-being and retirement are associated with higher Pleasantness. While individuals that do not rather report their occupation status showed negative association with Pleasantness. Further analysis revealed that psychological well-being was negatively associated with Eventfulness in men and individuals that did not report their occupation status. Additionally, we detected that Eventfulness is positively associated with unemployment, whereas it is negatively associated with gender (male) and retirement (Table 3).

The marginal and conditional R^2^ values are given in for each model in Table 3. In a mixed effects model, the marginal R^2^ represents the variance explained by the fixed effects (the individual-level independent variables) while the conditional R^2^ represents the variance explained by both the fixed and random effects (Nakagawa & Schielzeth, 2012). From the conditional R^2^, we can say that the full models explain 35.4% and 18.1% of the variance in Pleasantness and Eventfulness, respectively (Figure 1& 2). While the majority of the variance is explained by location-level differences (as confirmed by the intraclass correlation coefficients (ICC)), 1.4% of variance in Pleasantness and 3.9% of variance in Eventfulness is explained by the FOIs (i.e., psychological well-being and age) included as fixed effects.

**Figure 1 and 2.**
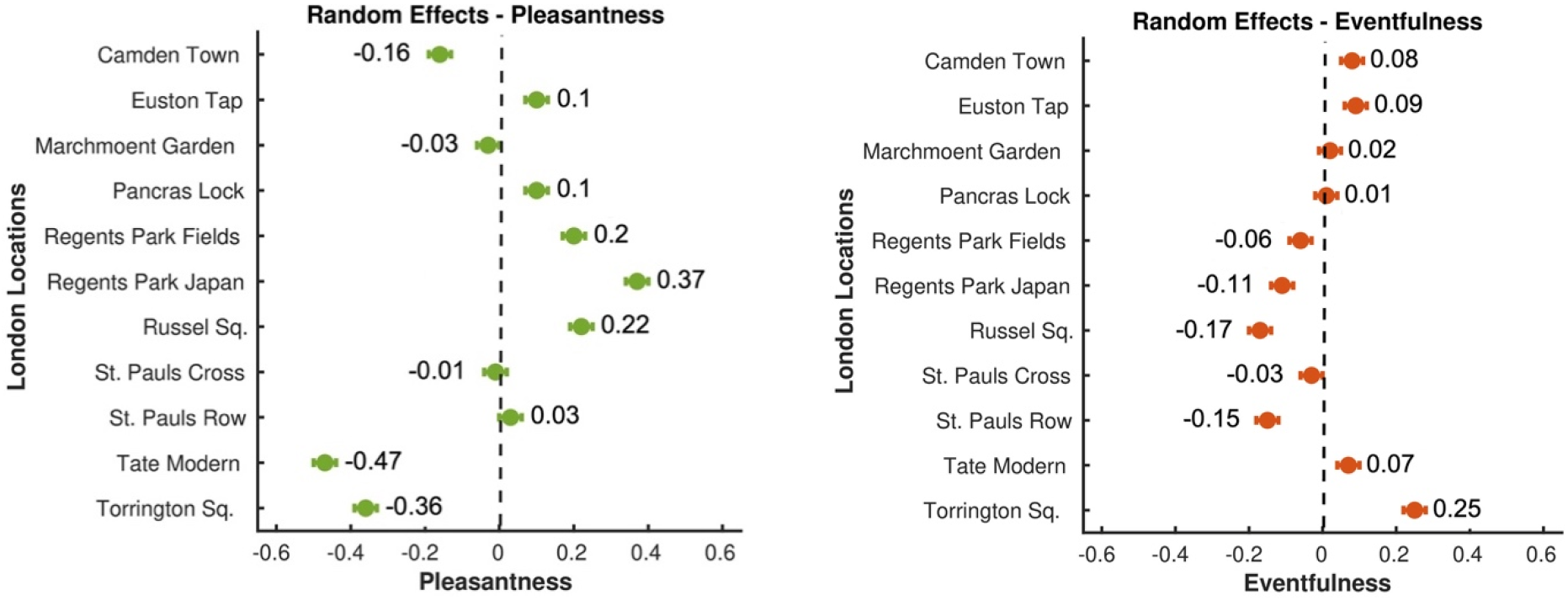
The summary result demonstrated in the random-effects figures gives the average from the distribution of Pleasantness (left) and Eventfulness (right) across locations.

## Discussion

For this study data of 1134 participants across 11 locations in London were included in the analysis. Our initial assumption was that an increased level of psychological well-being is associated with increased Pleasantness and Eventfulness assessments of the soundscape. Although the results showed that the psychological well-being was positively associated with Pleasantness, it was negatively associated with Eventfulness in males and individuals that did not report their occupations.

Then we hypothesized that differences in soundscape assessments are associated with demographic features. The results support this hypothesis to a certain degree. Occupation and gender appeared to be strong demographic factors influencing the Pleasantness and Eventfulness assessment. Retirement as occupation status showed to be positively attributed to the Pleasantness and negatively to the Eventfulness assessment. Further investigation revealed that the occupation (no occupation reported) was negatively associated with Pleasantness and gender (male) was negatively attributed to Eventfulness, whereas unemployment was positively associated with Eventfulness.

As expected, the majority of the total variance in the perceptual ratings is explained by the location-level differences (i.e., overall sound level) which represent primary contributing factors to the acoustic environment (see McDermott, 2012) and other non-acoustic factors. Approximately 3% of the variance is then explained by the combination of personal factors, which represent secondary contributing factors as defined by McDermott. Although the variance explained by these secondary factors is small compared to the primary factors, they are still found to contribute significantly. Furthermore, an additional 3 percentage points of explained variance would represent a meaningful improvement in the performance of predictive soundscape models based on in-situ measurements of varying soundscape types (Lionello, Aletta, & Kang, 2020) and should therefore be considered when constructing these models.

### Psychological well-being and its association with Pleasantness and Eventfulness

Our findings demonstrate a positive link between the perceived Pleasantness and participants’ psychological well-being, whereas the association between psychological well-being and Eventfulness is negative in males and individuals that did not report their occupations. Our results can be interpreted in light of previous research and it is consistent with the idea that psychological well-being underlies the perception of the external world (Kelley & Schmeichel, 2014) such as auditory input. While the enhanced global level of psychological state has a positive effect on auditory processing (Kumar, Sangamanatha, Vikas, 2013), there is evidence that suggests an impairment of early auditory processing (analysing, blending, and acoustic input segmentation) in individuals with poor psychological well-being (Kähkönen, Yamashita, Rytsälä, Suominen, Ahveninen, Isometsä, 2007). One of the potential trait biomarkers of poor psychological well-being such as depression (predominantly characterized by low mood and anhedonia (Erfanian, 2018) is the attenuation of neuronal activation in the auditory cortical area leading to alternations in auditory processing (Zwanzger, Zavorotnyy, Diemer, Ruland, Domschke, Christ et al., 2012).

### Demographic factors and their associations with Pleasantness and Eventfulness

#### Occupation status

According to our findings, occupation status, in particular ‘retirement’ and to a lesser degree, gender (male) were important factors in the pattern of soundscape assessments. It is worthwhile to highlight that ‘retirement’ factor can be potentially a proxy for age (>65) and gender (male). To explore the effect of occupation/retirement deeper on Pleasantness and Eventfulness we removed the occupation factor from the model. Age (*β* = 0.02, *p* = 0.05 for Pleasantness (*β* = -0.03, *p* = 0.01 for Eventfulness and gender (*β* = -0.04, *p* = 0.05 for Eventfulness then came out significant (see Appendix C). This would indicate that occupation status, particularly ‘retirement’, represents a group of older male individuals. Even though incorporation of occupation into our model complicates the interpretation of our outcome, it results in a slightly better fitting model (*R^2^_c_* for Pleasantness (0.354) and Eventfulness (0.181) relative to (0.345) for Pleasantness and (0.165) for Eventfulness in the model without occupation status which is why it is selected by the feature selection process. These findings are in line with previous research, suggesting significant differences among age groups in the soundscape of different acoustic environments (Ren, Kang, Liu, 2016; Yang & Kang, 2005). Our findings imply that an increase in age leads to an increase in the positive appraisal of the soundscape Pleasantness. This is supported by a study by Çakir Aydin & Yilmaz (2016) in which they found that soundscape pleasantness reported by young individuals was significantly lower than the other age groups. The results withstood a control for the effect of age on the soundscape’s pleasantness and eventfulness, suggesting that different neural and behavioural processes are responsible for the differences of soundscape appraisal in age.

One possibility is that age is associated with loss of function within the peripheral auditory system (hearing loss due to age or *presbycusis*) that may lead to the variation of the soundscape (Howarth and Shone, 2006). Higher tone frequencies have shown to be perceived less pleasant and more annoying relative to low tone frequencies (Landström, Kjellberg, SÖDerberg, Nordström, 1994) and age-related hearing loss is most marked at higher frequencies, so missing higher frequencies (that can be potentially unpleasant) may lead to an increase in soundscape pleasantness. Second, since the human brain is highly plastic throughout the life span, by ageing, the auditory processing changes due to the temporal coding of the auditory cortex (Bones & Plack, 2015; Babkoff & Fostick, 2017). Temporal coding is the ability of the brain to encode sensory information to the action potentials that rely on precise timing.

Last, age could potentially highlight the contextual role of the acoustic environment. Past experiences, memories, and even traumas give a particular context to our perception and shape the soundscape, making individual perception highly diverse, depending on the content of experience/memory. While the increase in age can lead to appreciating different sound elements, lower age seems to be related to more arousing and vibrant sounds (Yang & Kang, 2005).

Like age, gender was found to be associated with the soundscape Eventfulness. Past works have also reported that there are gender-related discrepancies in soundscape (Yang & Kang, 2005; Croome, 1977). These differences may be an indication of different auditory processing across genders. These differences are consistent with existing predictions of female top-down and male bottom-up strategies in spatial processing (ability to find where objects are in space) (Simon-Dack, Friesen & Teder-Sälejärvi, 2009).

### Soundscape Pleasantness and Eventfulness differences among locations

The Pleasantness and Eventfulness were significantly different among locations. The Pleasantness appeared to be highest in locations, dominating by nature sounds (i.e., Regents park Japan). In agreement with our results, Payne and colleagues (Payne, 2013) referred to the pleasantness dimension of the soundscape as the positive perception of natural places as well as the restorative capacity of the soundscape. Also, Zhang (2014) reported a significant impact of natural soundscape on individuals’ restorative experiences and boosting pleasantness. In the study by Axelsson et al. (2010) participants reported that the sound excerpts of natural components are more pleasant than human and technical sounds. Unlike Pleasantness, the Eventfulness increased the most in locations with dominant mechanical sounds (i.e., Euston Tap). These findings are supported by previous research done by Bradley & Lang (2000) and Hume & Ahtamad (2013). In both studies, unnatural and urban sound-clips (i.e., Fire engine siren and traffic noise), inherent in the traffic-dominant locations (i.e., Euston Tap) in our study, were rated highest in arousal and lowest in the pleasantness dimension. As formerly mentioned by Erfanian and colleagues (2019), throughout the soundscape literature, arousal has been applied as the equivalent of Eventfulness and indicated on the Y-axis of the circumplex model (Erfanian et al, 2019; Axelsson et al., 2010).

These results insinuate the notion that there are multiple primary factors (McDermott, 2012) that contribute to the perception of the acoustic environment which should be considered important by urban designers and policymakers. It is expected that understanding these factors will provide multidimensional knowledge in guiding the implementation of the technological the infrastructure of smart cities.

## Conclusion

We conducted a linear mixed-effects model to show the associations of psychological well-being, demographic factors with the soundscape Pleasantness and Eventfulness. The findings indicate that psychological well-being is positively associated with Pleasantness and negatively associated with Eventfulness in males and individuals that did not report their occupations. We further demonstrated that the occupation status, in particular retirement as a proxy of age and gender, was attributed to Pleasantness and Eventfulness. The findings of this study offer empirical grounds for developing and advancing theories on the influence of psychological well-being and demographic characteristics on the perception of the acoustic environment namely the soundscape.

## Appendix A

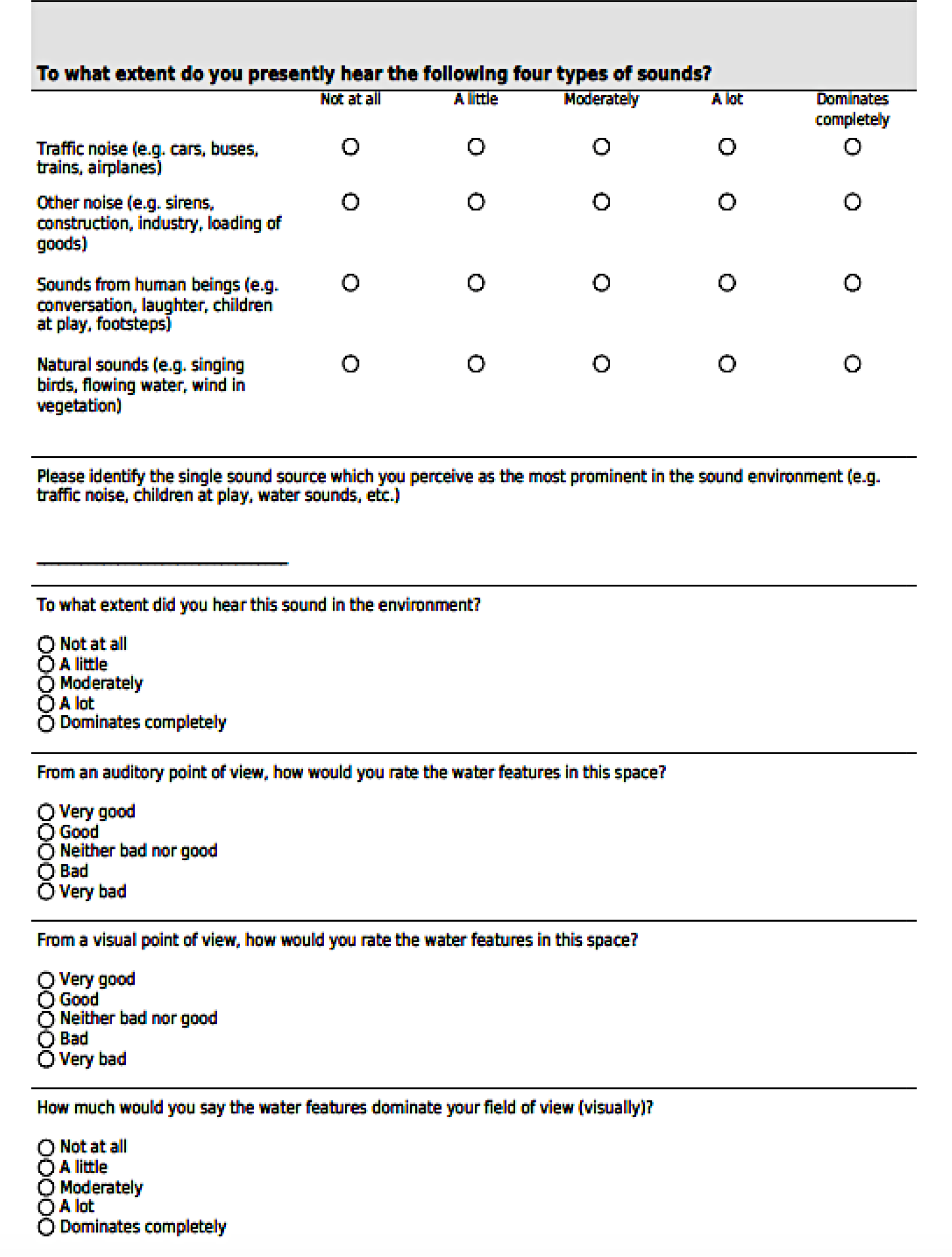

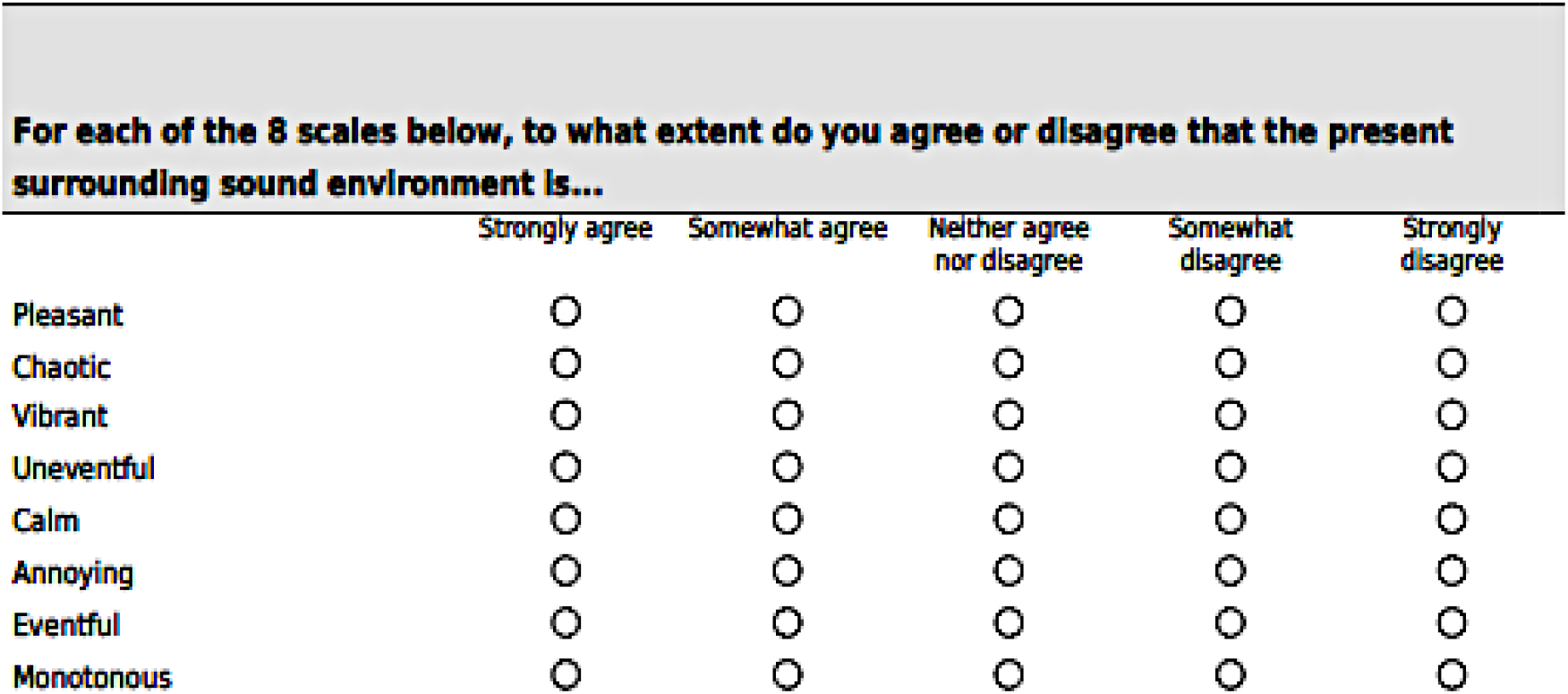

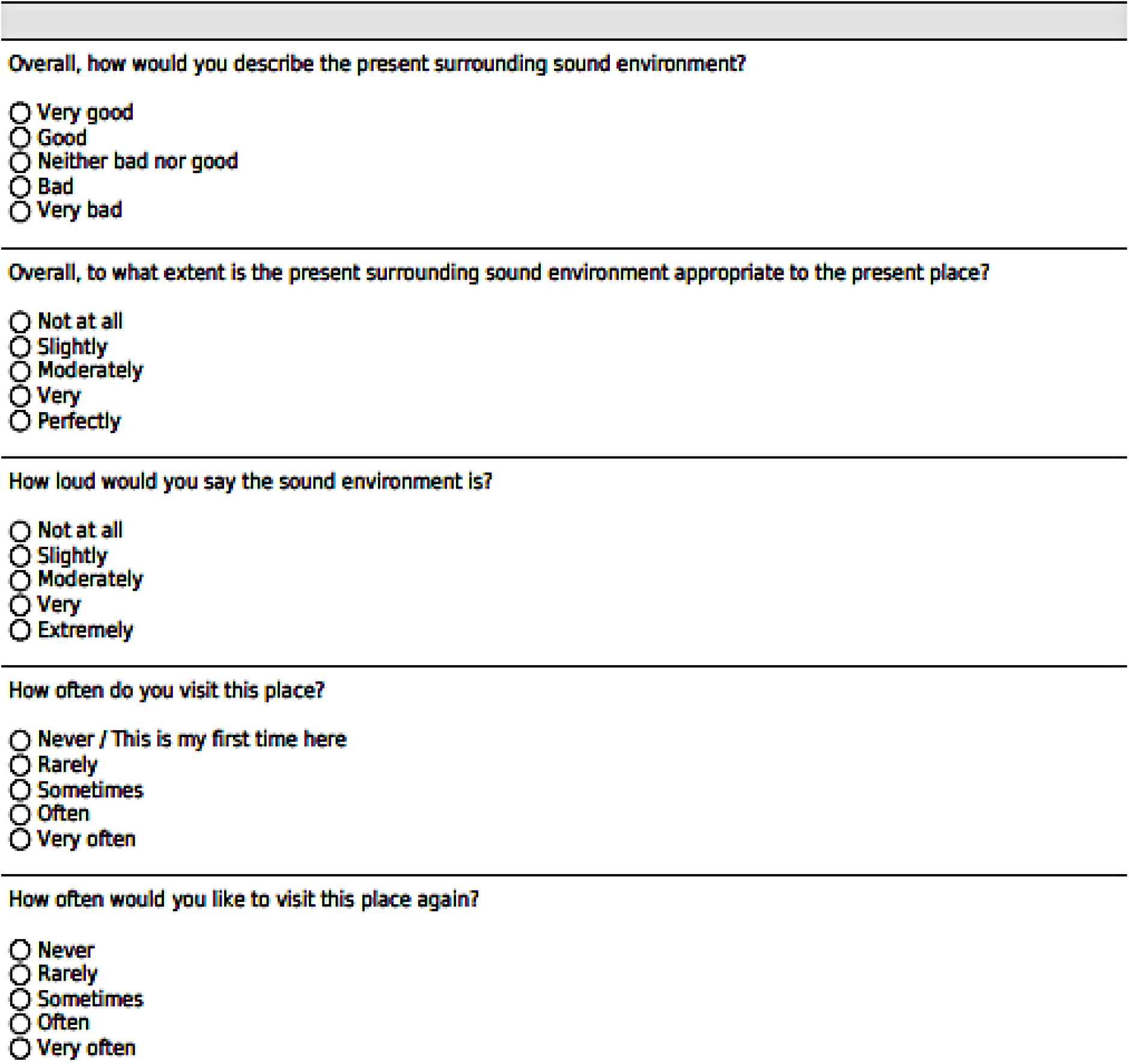

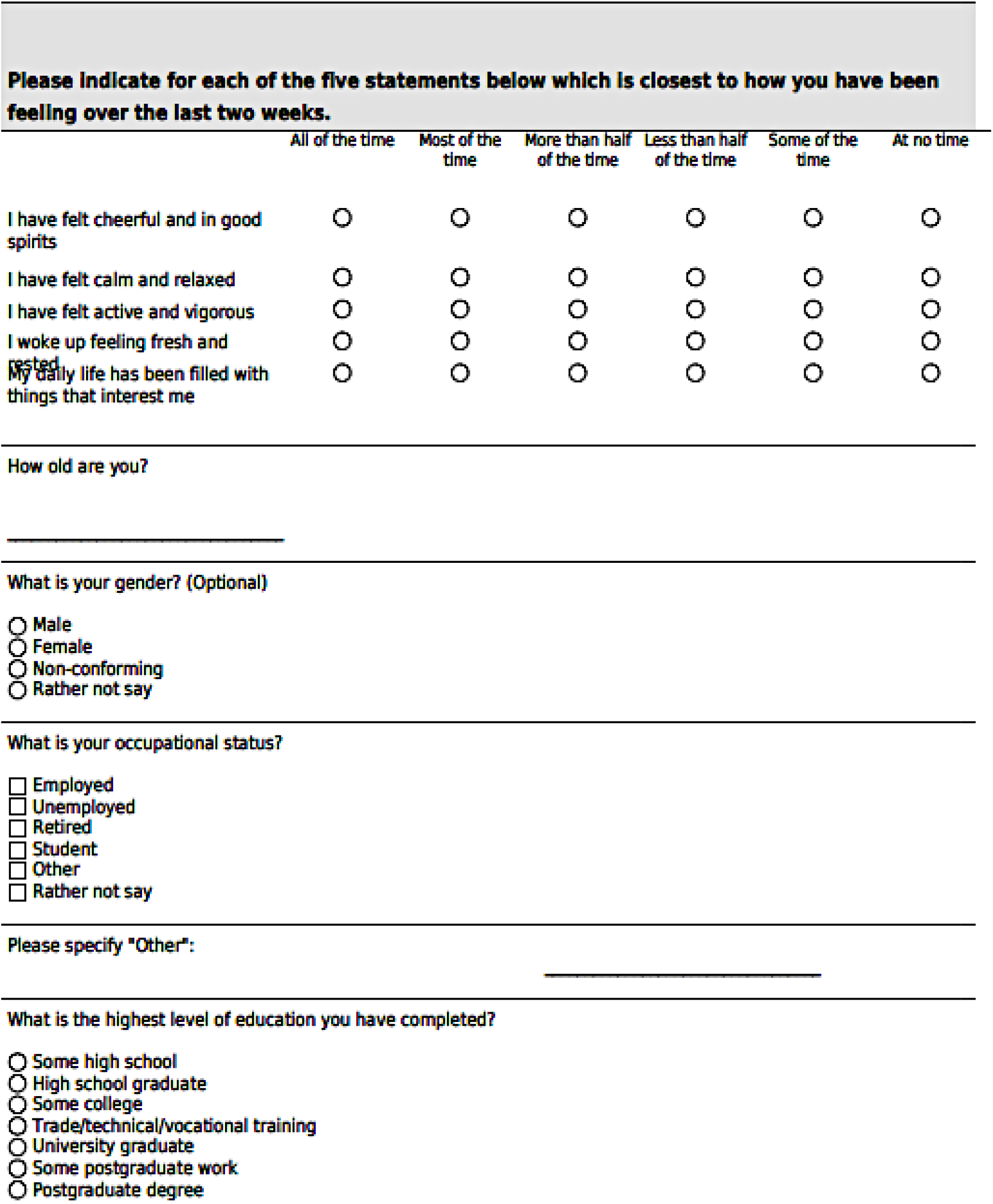

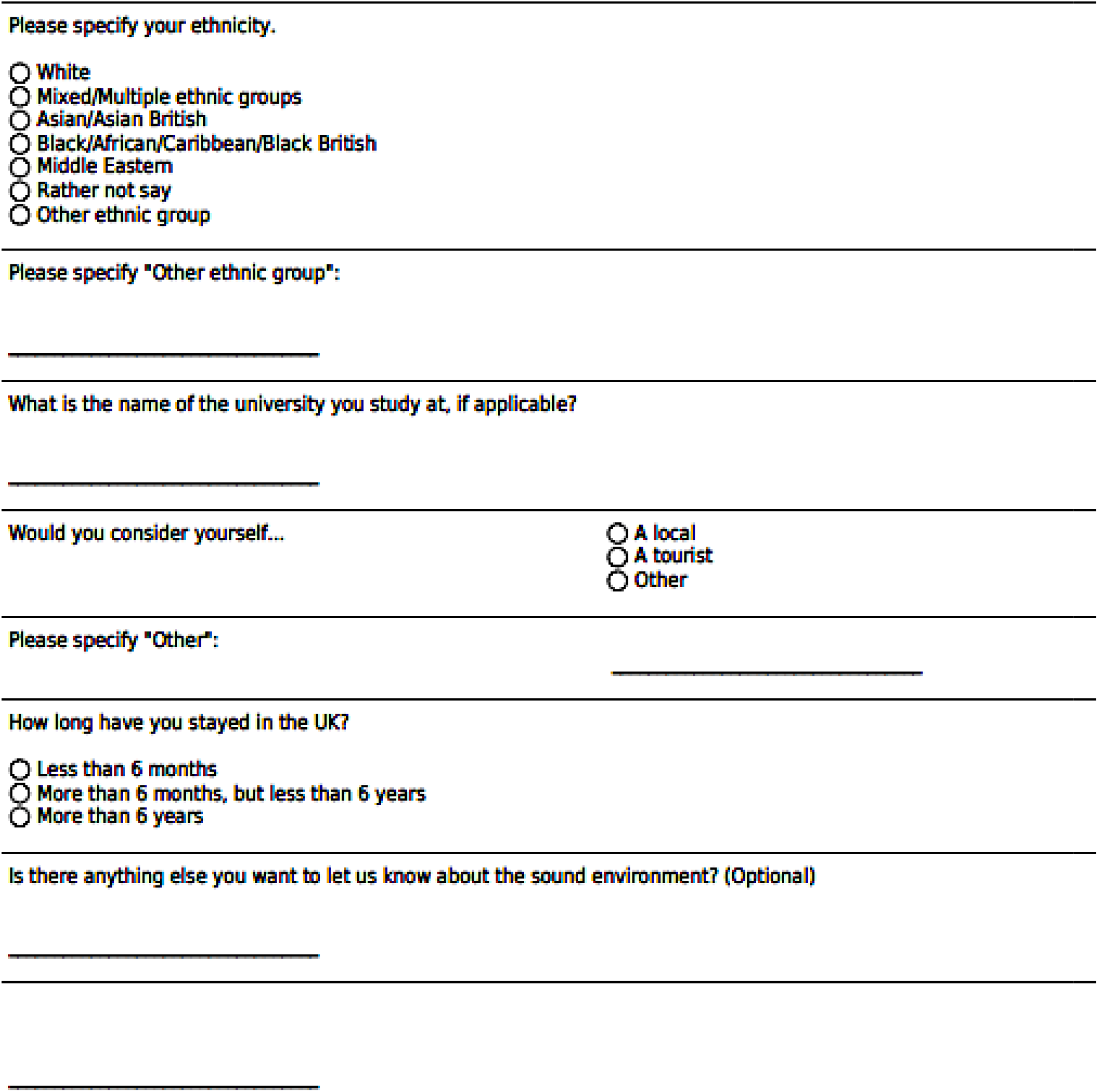

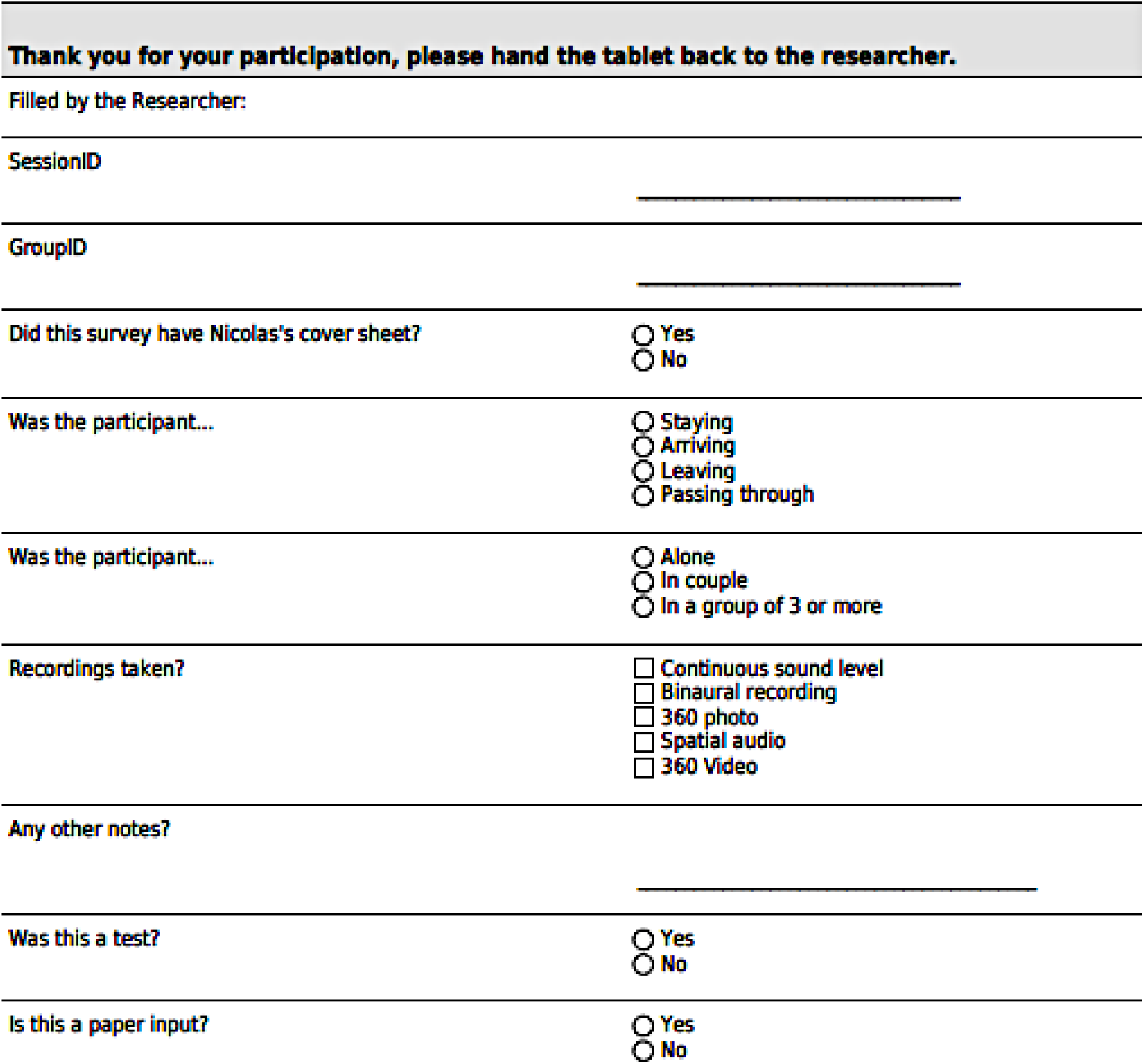

## Appendix B

**Table B.1.** depicts the minimum and maximum value of acoustic metrics of each location during the survey periods.

| Location | L<sub>Aeq</sub> | L<sub>A90</sub> | L<sub>A10</sub> | L<sub>A10</sub>- L<sub>A90</sub> | L<sub>AFmax</sub> | L<sub>AFmin</sub> |
| --- | --- | --- | --- | --- | --- | --- |
| Camden Town | 69- 84 | 62-72 | 70-90 | 7-25 | 92-100 | 55-62 |
| Marchmont Garden | 56-58 | 48-51 | 57-62 | 7-12 | 83-94 | 45-46 |
| Pancras Lock | 59-61 | 55-56 | 62-63 | 7 | 87-104 | 49-50 |
| Regents Park Fields | 53-64 | 45-46 | 55-61 | 9-16 | 82-88 | 42-44 |
| Regents Park Japan | 62 | 60 | 62 | 2 | 83 | 57 |
| Russell Square | 66-73 | 64-72 | 69-74 | 2-5 | 87-95 | 59-68 |
| Tate Modern | 62-63 | 55-58 | 64-65 | 8-9 | 85-88 | 51-53 |
| Torrington Square | 64-68 | 57-58 | 66-67 | 9 | 92-106 | 51 |
| St. Paul's Cross | 61 | 56 | 62 | 6 | 84 | 53 |
| St. Paul's Row | 62 | 59 | 64 | 6 | 81 | 55 |
| Euston Tap | 69-73 | 63-64 | 70-73 | 7-10 | 92-104 | 58-60 |

**Table B.2.** demonstrates the sound source composition of the selected locations in London.

| Locations | N | Natural | Traffic | Human | Other |
| --- | --- | --- | --- | --- | --- |
| Camden Town | 107 | 1.33 | 3.75 | 3.26 | 2.66 |
| Euston Tap | 102 | 1.66 | 3.71 | 2.56 | 2.95 |
| Marchmont Garden | 106 | 2.59 | 2.65 | 2.66 | 2.45 |
| Pancras Lock | 99 | 2.38 | 2.43 | 2.48 | 3.28 |
| Regents Park Fields | 116 | 3.09 | 2.4 | 2.9 | 1.87 |
| Regents Park Japan | 93 | 4.02 | 1.88 | 2.53 | 1.52 |
| Russell Square | 149 | 3.27 | 2.77 | 3.04 | 2.16 |
| St. Pauls Cross | 66 | 2.3 | 2.57 | 3.31 | 2.1 |
| St. Pauls Row | 69 | 1.76 | 2.55 | 3.45 | 2.25 |
| Tate Modern | 156 | 2.58 | 2.5 | 3.64 | 2.14 |
| Torrington Square | 117 | 1.93 | 3.19 | 3.25 | 2.81 |

## Appendix C

**Table C1.**
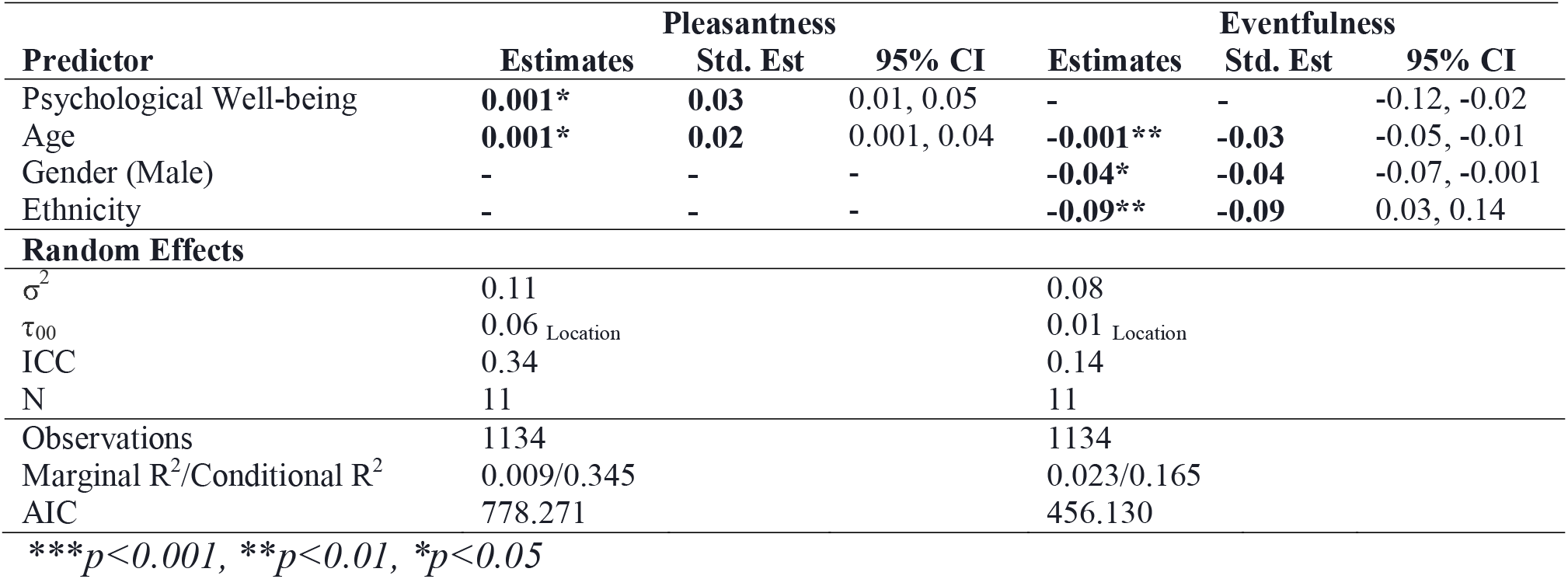
Fixed and random effects in a linear mixed model explaining variation in the soundscape Pleasantness and Eventfulness while controlling for psychological well-being and demographic factors, excluding occupation.

## Appendix D

**Figure D1a.**
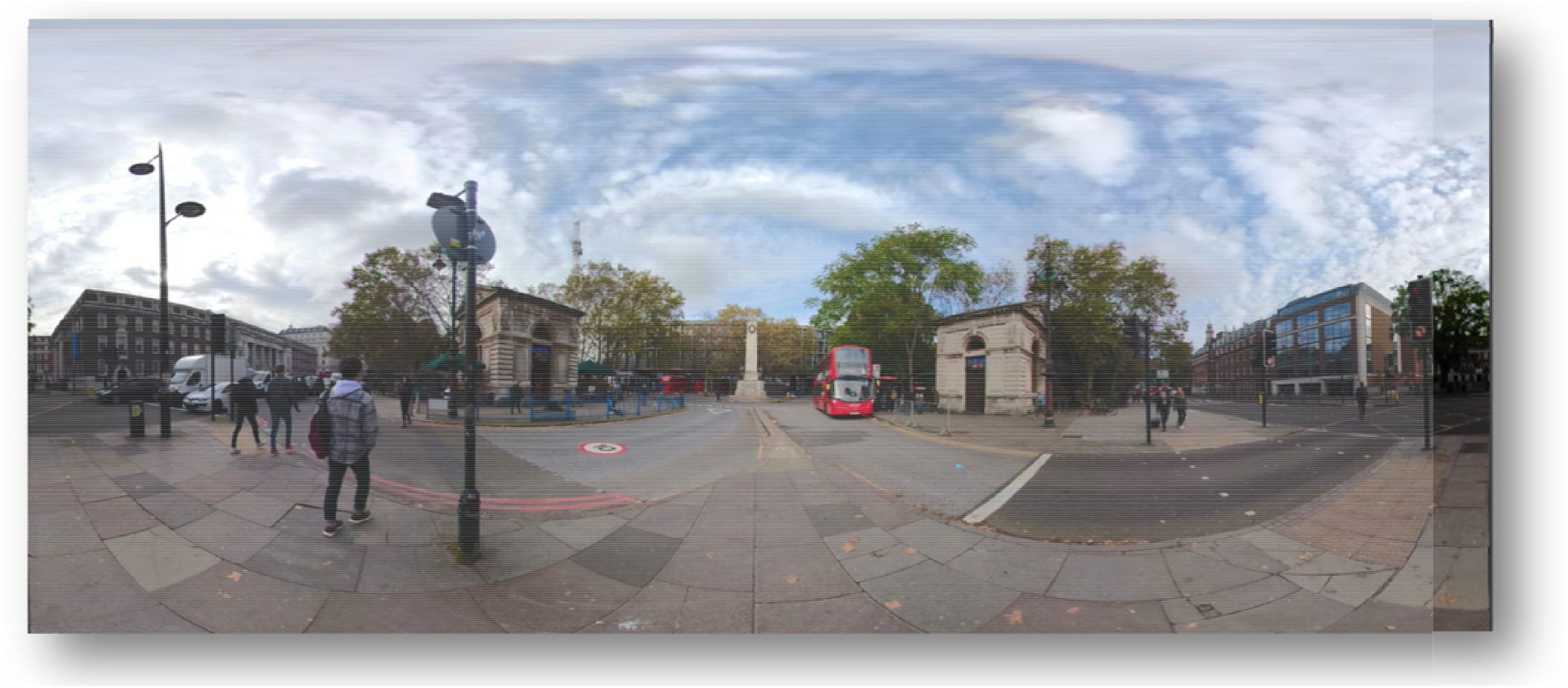
shows Euston Tap in London represents an acoustic environment dominated by traffic noise.

**Figure D1b.**
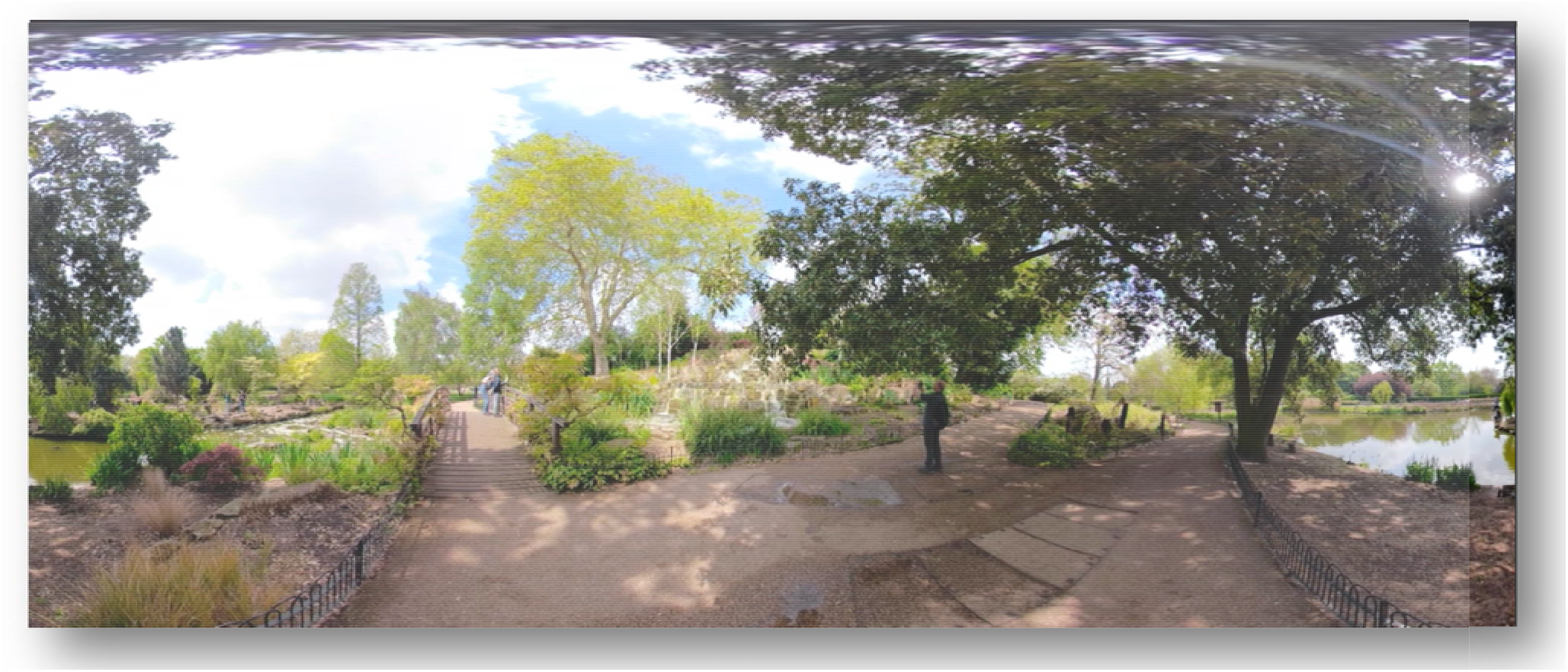
shows Regents Park Japan in London represents an acoustic environment with natural environmental sound

**Figure D1c.**
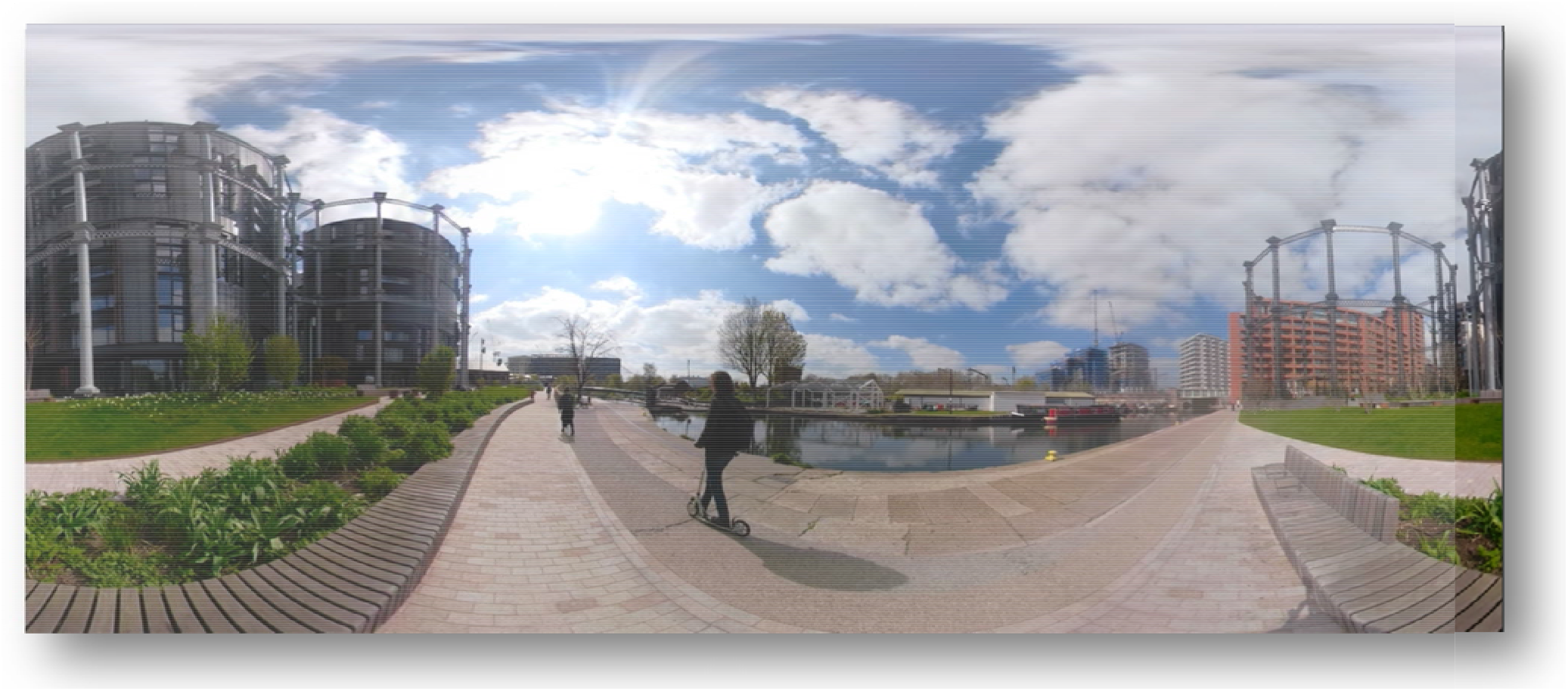
*shows* Pancras Lock in London represents an acoustic environment with a mix of natural and unnatural environmental sound

## Footnotes

1 International Organization for Standardization/Technical Specification (12913-3, 2019) deals with work still under technical progress/development, or where it is believed that there will be a future, but not immediate, possibility of agreement on an International Standard. A Technical Specification is published for immediate use, but it also provides a means to obtain feedback.

2 The ISO/TS 12913-2:2018 specifies requirements and provides supporting information on data collection and reporting for soundscape studies, investigations and applications.

3 The ISO/TS 12913-3:2019 provides requirements and supporting information on analysis of data collected in-situ.

## Notes

### Competing Interest Statement

The authors have declared no competing interest.

